# Decoding Radiation-Induced Transcriptomic Signatures of Human Leukocytes using Long-Read RNA-Seq: Clinical and Biodosimetric Implications

**DOI:** 10.64898/2026.02.04.703787

**Authors:** Ahmed Salah, Heinz Schmidberger, Federico Marini, Sebastian Zahnreich

## Abstract

**Background:** Gene expression profiling in radiation-exposed blood is a valuable tool for biodosimetry and clinical research. Evaluating the blood’s transcriptomic radiation response provides insight into absorbed dose, hematotoxicities, and immune reactions. However, detailed analysis using long-read RNA sequencing is currently limited in its diffusion, despite the potential additional insights that could be extracted, including novel isoform discovery and on-the-field gene expression studies, owing to its portability.

**Results:** In this study, we utilized Oxford Nanopore Technologies’ long-read RNA sequencing on human peripheral blood leukocytes from three healthy donors 6 hours after exposure of whole peripheral blood to 4 Gy of X-rays. Compared to sham-irradiated (0 Gy) blood samples, gene-level differential expression analysis identified 117 upregulated and 66 downregulated genes, including canonical DNA damage repair and inflammatory responses. At the transcript level, 102 transcripts were significantly upregulated, and 17 were downregulated, revealing isoform-specific regulation that was not captured at the gene level. Notably, *IL32*, which was not detected as significantly changed at the gene level, exhibited significant upregulation of two transcript isoforms (log2 fold-changes of 1.19 and 1.99), while *WDR74*, *ITM2B*, *AK2*, and *RPS19* displayed changes in transcript usage following irradiation. Leveraging long-read RNA sequencing, we further identified novel candidate isoforms, including protein-coding isoforms of *PHPT1* and *FBXO22* as well as non-coding candidates. Notably, the *FBXO22* isoform showed evidence of altered protein domain architecture relative to the reference isoform.

**Conclusions:** This study is the first to systematically investigate differential transcript usage in the leukocyte fraction of irradiated human whole blood using long-read RNA sequencing. We confirmed known radiation-response signatures at the gene level and identified isoform-level transcriptomic changes, including several isoforms that were undetectable in our previously generated matched short-read dataset. These findings demonstrate the potential of long-read RNA sequencing to provide a more detailed view of radiation-induced transcriptomic alterations, with potential applications in biodosimetry and clinical settings.

## Introduction

Exposure to ionizing radiation (IR) arises from natural background sources, medical procedures, occupational settings, or accidental incidents [1]. In all cases, blood is unavoidably irradiated, causing hematotoxic and immune effects while also serving as a key tissue for biological dosimetry [2, 3].

IR induces various types of DNA lesions, with DNA double-strand breaks (DSBs) representing the most critical damage [4–7]. High-linear energy transfer (LET) radiation, such as neutrons, alpha particles, and heavy ions, causes more complex DNA damage than low-LET radiation (e.g., photons) due to its denser spatial energy distribution, resulting in higher relative biological effectiveness (RBE) and greater health risks [1, 8, 9]. The quantitative and qualitative detection of IR-induced DSBs, either directly or through non-repair or misrepair, underpins traditional biodosimetric assays in peripheral blood lymphocytes (PBLs). These include chromosome aberration assays, the micronucleus assay, and immunohistochemical detection of DSB repair proteins such as γH2AX, 53BP1, or RPA to reconstruct absorbed doses and exposure scenarios [10–12]. However, DNA damage also strongly triggers broader cellular and transcriptomic responses [13]. Consequently, gene expression analysis in whole blood or isolated PBLs exposed to IR has become an established approach in recent years, both in biodosimetry and in the clinical context of radiotherapy (RT) [14–16].

IR also strongly modulates the hematological immune system, showing anti-inflammatory effects at low doses (≤0.5 Gy) and pro-inflammatory effects at high doses (≥2 Gy), in addition to DNA damage and cell death signaling [17]. Such dose-dependent immune responses to IR are particularly relevant clinically, e.g., during RT, where IR is applied at low doses to treat benign inflammatory diseases and at high doses to treat malignancies [18–21]. Therefore, the immunomodulatory and adverse hematologic effects of RT are gaining high importance, particularly with its growing combination with immuno-oncological treatments [22, 23].

A key advantage of gene expression analysis via transcriptomics over traditional, time-consuming, and labor-intensive biodosimetric assays is its rapid, scalable, and cost-effective capability of measuring all the genes at once for assessing IR exposure and responses. However, both general and IR-induced gene expression are strongly influenced by confounding factors, such as age, gender, and ethnicity, which must be considered before concluding [24, 25]. Therefore, a deeper understanding of IR-induced gene expression and related molecular pathways is necessary for both biodosimetry and clinical applications, including side effect and toxicity prediction, and treatment planning in multimodal settings such as radioimmunotherapy.

Gene expression profiling shows a strong correlation with absorbed radiation dose, both in *ex vivo* and *in vivo* irradiated blood samples [26–31], and specific gene signatures can predict radiation exposure status and dose level with over 90% accuracy in both mice and humans [32, 33]. A master regulator of the DNA damage response (DDR) is p53, which transcriptionally controls several IR-responsive genes, as listed in reference data for biodosimetry, including *FDXR*, *PCNA*, *BRCA1*, *CDKN1A*, *MDM2*, and *BBC3* [34–36]. In biodosimetry, however, there remains an urgent need to develop specific gene expression signatures to predict absorbed dose and, ideally, to distinguish between different radiation qualities with varying RBE. Most previous biodosimetry studies based on gene expression have relied on microarray or quantitative PCR (qPCR) platforms, which, while effective for targeted analyses, provide limited depth and coverage of the transcribed mRNA population compared to RNA sequencing (RNA-seq) [37, 38]. RNA-seq has emerged as a powerful tool to address these gaps, enabling genome-wide quantification of transcript expression and regulatory changes with high sensitivity and dynamic range [39]. However, only a limited number of studies have leveraged genome-wide transcriptomics by short-read (SR) RNA-seq to investigate IR-related transcriptional responses [17, 40]. To enable the development of reproducible and IR-specific gene signatures, more high-throughput gene expression and meta-analysis studies are highly warranted.

Recently, long-read (LR) RNA-seq, also referred to as third-generation RNA-seq, has expanded the scope of transcriptomic profiling beyond gene-level expression to isoform-level resolution [41]. This technology captures full-length transcripts in a single read, allowing the accurate characterization of alternative splicing and novel isoform usage that may arise in response to IR [42–44]. Due to its affordability and portability, Oxford Nanopore Technologies (ONT) provides a valuable in-the-field platform for gene expression studies.

In the present study, we employed ONT LR RNA-seq to profile the transcriptomic response of the total leukocyte fraction of peripheral blood from three healthy donors 6 h after *ex vivo* exposure to 4 Gy X-rays. Our results not only reproduce known radiosensitive genes but, more importantly, extend current understanding by identifying new isoforms in IR-sensitive genes.

## Methods

### Blood sampling, irradiation, culturing, and RNA isolation

Nine ml of whole blood was collected from three healthy donors (two males and one female) by venipuncture into Ethylenediaminetetraacetic acid (EDTA) tubes (S-Monovette® EDTA K3E, Sarstedt, Nuembrecht, Germany). Ethical approval was obtained from the Ethics Committee of the Rhineland-Palatinate Chamber of Physicians [No. 2023-17191], and all research was performed following relevant guidelines and regulations. All donors provided informed consent, and research was performed in accordance with the Declaration of Helsinki. All three donors were non-smokers, and their ages at the time of blood collection were 46 (male), 31 (male), and 29 (female) years, respectively. Samples were exposed to 4 Gy X-rays at room temperature, with sham-irradiated (0 Gy) controls maintained under identical conditions in the radiation device control room, as described in detail previously [17]. After irradiation, whole blood was incubated in EDTA tubes in a humidified atmosphere at 37 °C and 5% CO_2_ for 6 h. RNA was extracted from total leukocytes using the LeukoLOCK Total RNA Isolation System (ThermoFisher Scientific) according to the manufacturer’s protocol. Globin and ribosomal RNA (rRNA) fractions can be found in Additional File 1.

### RNA sequencing

RNA concentration was quantified using the Qubit RNA High Sensitivity Assay (Thermo Fisher Scientific). RNA integrity was assessed as RNA quality number (RQN) using a Fragment Analyzer 5200 with the DNF-471 RNA kit (Agilent Technologies). RQN values are provided in Table 1.

**Table 1.**
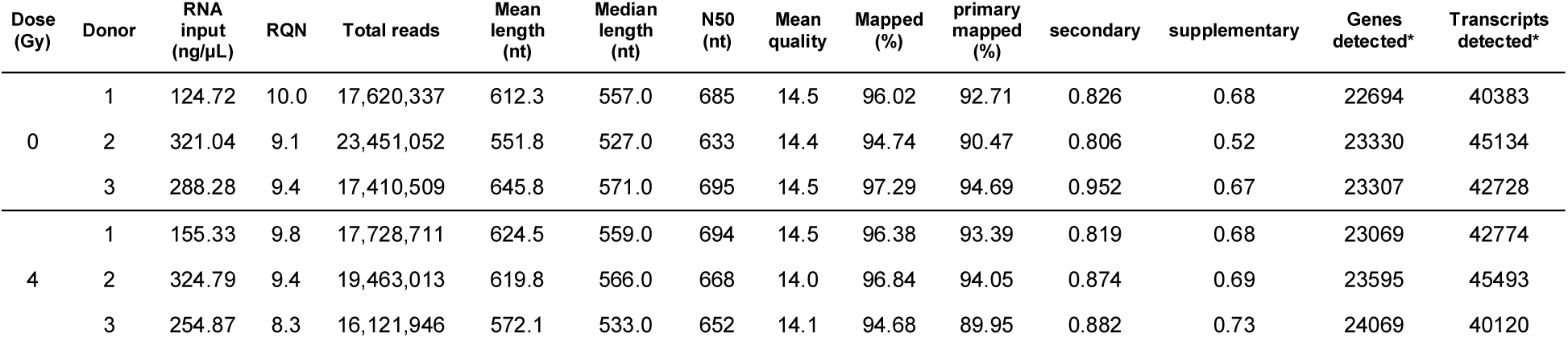
Per-library sequencing, alignment, and quantification metrics for the LR RNA-seq libraries. Genes and transcripts detected reflect counts from IsoQuant’s gene- and transcript-level grouped count outputs prior to low-expression filtering; filterByExpr-based filtering (edgeR) was subsequently applied jointly across all libraries for differential expression analysis (13,020 genes; 14,146 transcripts retained after filtering). IsoQuant read assignment (pooled across all six libraries, as IsoQuant was run jointly on all samples with no per-library assignment tracking available): of 113,225,145 total assigned reads, 22.0% were classified as unique, 12.2% as unique with minor difference, 53.4% as ambiguous, 18.6% as inconsistent (combined subtypes), 0.1% as intergenic, and 0.9% as noninformative. RQN, RNA quality number; nt, nucleotides.

Libraries were prepared using the PCR-cDNA Barcoding Kit SQK-PCB114.24 (ONT) according to the manufacturer’s instructions. Briefly, 750 ng per sample total RNA was used for reverse transcription and strand-switching with Maxima H Minus Reverse Transcriptase (Thermo Fisher Scientific). The resulting cDNA was amplified using LongAmp® Hot Start Taq Master Mix (New England Biolabs) for 14 PCR cycles following the ONT protocol. Library concentration was measured using the Qubit dsDNA assay (Thermo Fisher Scientific). Six barcoded libraries were pooled equimolarly (20 fmol per library). Rapid adapters were ligated to the pooled cDNA, and the library was loaded onto an R10.4.1 PromethION flow cell (FLO-PRO114M). Sequencing was performed on a PromethION 2 Solo for 72 h with FAST basecalling and barcode balancing enabled.

For downstream analysis, super-accuracy (SUP) basecalling was performed using Dorado basecaller v0.8.2 (https://github.com/nanoporetech/dorado) with model dna_r10.4.1_e8.2_400bps_sup@v5.0.0. Adapter trimming and barcode demultiplexing were carried out using dorado trim and dorado demux, respectively.

### Processing of RNA-seq data

LR sequencing data were processed as follows. Base-called reads were quality-assessed with NanoPlot (version 1.44.0) [45] and aligned to the GRCh38 using minimap2 (version 2.28-r1209) [46] in spliced alignment mode for ONT cDNA reads (-ax splice -uf -k14), with resulting alignments coordinate-sorted and indexed using samtools [47]. Mapping rates were between 94.68–97.29% across all the libraries. Transcript and gene quantification, together with novel transcript model construction, were performed using IsoQuant (version 3.6.2) against the GENCODE v47 annotation (gencode.v47.annotation.gtf) supplied as the reference gene database (--genedb), run in default ONT model-construction mode (-- model_construction_strategy default_ont) with the --complete_genedb and --report_novel_unspliced true options enabled [48]. This configuration runs IsoQuant in a combined reference-guided and transcript-discovery mode: reads consistent with annotated GENCODE v47 transcripts are quantified against that reference, while reads not fully explained by the existing annotation are used to construct and quantify novel, unannotated transcript models. Gene- and transcript-level count matrices were obtained directly from IsoQuant’s grouped count outputs, each spanning all six libraries and including both annotated and unannotated features.

Before differential expression analysis (DEA), low-abundance genes and transcripts were filtered out independently from the gene- and transcript-level count matrices using the filterByExpr() function from edgeR (version 4.7.5) [49], yielding 13,020 genes and 14,146 transcripts retained for gene- and transcript-level DEA, respectively.

### Differential expression and differential transcript usage analysis

DEA was conducted using the edgeR package (version 4.7.5) [49], with the false discovery rate (FDR) cutoff set to 0.05. Sham-irradiated (0 Gy) samples were used as the reference level. The model specification (∼Donor + Treatment) was set to account for the dose effects and inter-donor variability. Genes were considered differentially expressed if their adjusted p-value (p_adj_) (Benjamini-Hochberg procedure; [50]) was equal to or smaller than 0.05. We also applied the DESeq2 R package (version 1.49.4) for statistical analysis of quantified reads using the same design settings and cutoffs [51].

A leave-one-donor-out sensitivity analysis was performed by repeating the gene-level edgeR differential expression pipeline three times, each time excluding one donor (n = 2 per group), and comparing the resulting log fold change (LFC) estimates and significant gene sets to those from the full three-donor model.

Sequencing-depth saturation was assessed using the subSeq Bioconductor package (version 1.40.0) [52] by repeatedly subsampling the filtered raw count matrix to 5%, 10%, 20%, 30%, 40%, 50%, 60%, 70%, 80%, 90%, and 100% of the observed sequencing depth. At each subsampling level, 20 independent subsampling replicates were generated, and differential expression was re-estimated using the same donor-adjusted edgeR quasi-likelihood model as used for the primary analysis (∼Donor + Treatment). Differentially expressed genes (DEGs) were defined using Benjamini–Hochberg p_adj_ ≤ 0.05, consistent with the primary DEA. The full-depth analysis (100% of the observed sequencing depth) was used as the reference. Saturation was evaluated by quantifying the number and proportion of full-depth DEGs recovered at each sequencing depth, together with the Spearman correlation and concordance of estimated LFCs relative to the full-depth analysis. This analysis was used to assess the extent to which reduced sequencing depth affected DEG detection and the stability of estimated transcriptional responses.

To enable direct comparison with the LR dataset, SR RNA-seq data from our previously published study of X-irradiated human blood [17] were re-analyzed as follows: SR reads, 150 nucleotides, generated from irradiated whole blood cultured *ex vivo*, with RNA extracted directly from whole blood followed by globin and rRNA depletion, were quantified at the transcript level using Salmon (Version 1.10.3) [53] and imported into R using tximport (version 1.22.1) [54], against the GENCODE v46 reference annotation. For gene-detection comparisons, the SR and LR gene-level count matrices were restricted to their shared gene universe (genes annotated in both GENCODE v46 and v47), and a common low-expression filter (filterByExpr, edgeR) was applied independently to each platform to define detected genes. For differential expression comparisons, the SR dataset was further restricted to the same two dose groups and time point used in the LR study (0 Gy and 4 Gy, 6 h post-irradiation, n = 3 donors per group), rather than the full dose-response series of 0.5-4 Gy used in the original publication, to ensure a comparable experimental design between platforms. Differential expression in this restricted SR comparison was assessed using DESeq2 with independent filtering, consistent with the original publication’s methodology; differential expression in the LR dataset was assessed using edgeR with filterByExpr-based filtering, as described above.

Directional concordance between SR and LR LFC was assessed using a two-sided exact binomial test against a null concordance rate of 0.5. Gene-set-level enrichment of the same signature within the LR differential expression model was additionally assessed using a competitive gene-set test (mroast, limma; (Version 3.66.0) [55] design: ∼Donor + Treatment; contrast: 4 Gy vs. 0 Gy), evaluating the mixed-direction null hypotheses.

Differential transcript usage (DTU) was performed using the diffSplice pipeline, available in the edgeR package [56, 57]. FDR was corrected using the stageR workflow (version 1.31.0) [58].

Structural quality control of the candidate unannotated transcripts was performed using SQANTI3 (v6.0.2) [59], based on LR data and the GENCODE v47 reference annotation and genome, incorporating SR junction coverage from STAR-aligned RNA-seq data (SJ.out.tab files) (version 2.7.11b) [60] available for the same donors. Open reading frame (ORF) prediction and coding-potential scoring were performed using SQANTI3’s integrated psauron module. The standard SQANTI3 rules-based filter (default parameters) was applied to flag transcripts with structural features associated with technical artefacts (e.g., non-canonical splice junctions, RT-switching, or absent coverage support). Results were cross-referenced against each transcript’s independent GffCompare classification as a second layer of structural evidence.

For each of the filter-passed transcripts, SR junction support was independently re-derived from STAR SJ.out.tab files, and LR support was assessed by directly scanning ONT alignment Compact Idiosyncratic Gapped Alignment Report (CIGAR) strings for a spliced (’N’ operation) gap matching each junction’s exact genomic coordinates (±2 bp tolerance), implemented in a custom Python/pysam script. For each junction this provided an independent SR read count, a locus-wide SR activity check, and the number of LR libraries and reads independently confirming the junction, summarized across all six sequencing libraries.

### Functional enrichment analysis

Functional enrichment analysis was performed using clusterProfiler (Gene Ontology (GO) over-representation and Kyoto Encyclopedia of Genes and Genomes (KEGG) pathway analysis), and the Gene Set Enrichment Analysis (GSEA) implementation in clusterProfiler/MSigDB (GSEA using MSigDB C5 BP gene sets), each applied to the differentially expressed gene list described above with an explicitly defined background universe [61, 62]. The enrichment results were visualized and summarized using the GeneTonic package (version 3.3.3) [63]. Enriched pathways were displayed as heatmaps, with color-coded standardized Z-scores for expression values. These values were obtained after a regularized logarithm transformation, facilitating comparison across samples.

### Identification of protein-coding transcripts

Candidate ORFs were identified using TransDecoder (version 5.7.1) (https://github.com/TransDecoder/TransDecoder): TransDecoder.LongOrfs (minimum ORF length: 100 amino acids), with ORFs scored using TransDecoder.Predic (default parameters); ORFs with a positive coding-likelihood score (log-likelihood ratio > 0) were retained. Novel transcript isoform classification was performed using GffCompare (version 0.12.10) [64]. Coding potential prediction was performed using the Coding Potential Calculator 2 (CPC 2.0) online tool [65] (https://cpc2.gao-lab.org/, access date Oct 29, 2025) using coding probability score threshold of 0.5 and the Coding-Potential Assessment Tool (CPAT) (version 3.0.5) [66].

Predicted ORFs for the coding-candidate transcripts were searched against the NCBI non-redundant (nr) protein database using BLASTp, restricted to Homo sapiens, with a maximum of 5 target sequences reported per query [67].

Functional domain content of the predicted ORFs for the coding-candidate transcripts was assessed using InterProScan [68] (v5.78-109.0) accessed via the EBI InterProScan web service (https://www.ebi.ac.uk/interpro/) on 21 July 2026, run with default parameters against the full set of member databases included in the standard InterProScan analysis (including Pfam, PANTHER, and SUPERFAMILY). Domain architecture of each candidate ORF was compared against that of the corresponding GENCODE v47 reference protein, analyzed under identical InterProScan parameters.

## Results

### Gene-level ONT RNA-Seq data reveal radiation-induced DNA damage and immune signaling in human leukocytes

Whole-blood samples obtained from three healthy donors were sham-irradiated (0 Gy) or exposed to 4 Gy of X-rays and subsequently incubated at 37 °C for 6 h. The workflow of this study is illustrated in Figure 1. Sequencing and alignment quality metrics for all six libraries are summarized in Table 1.

**Figure 1:**
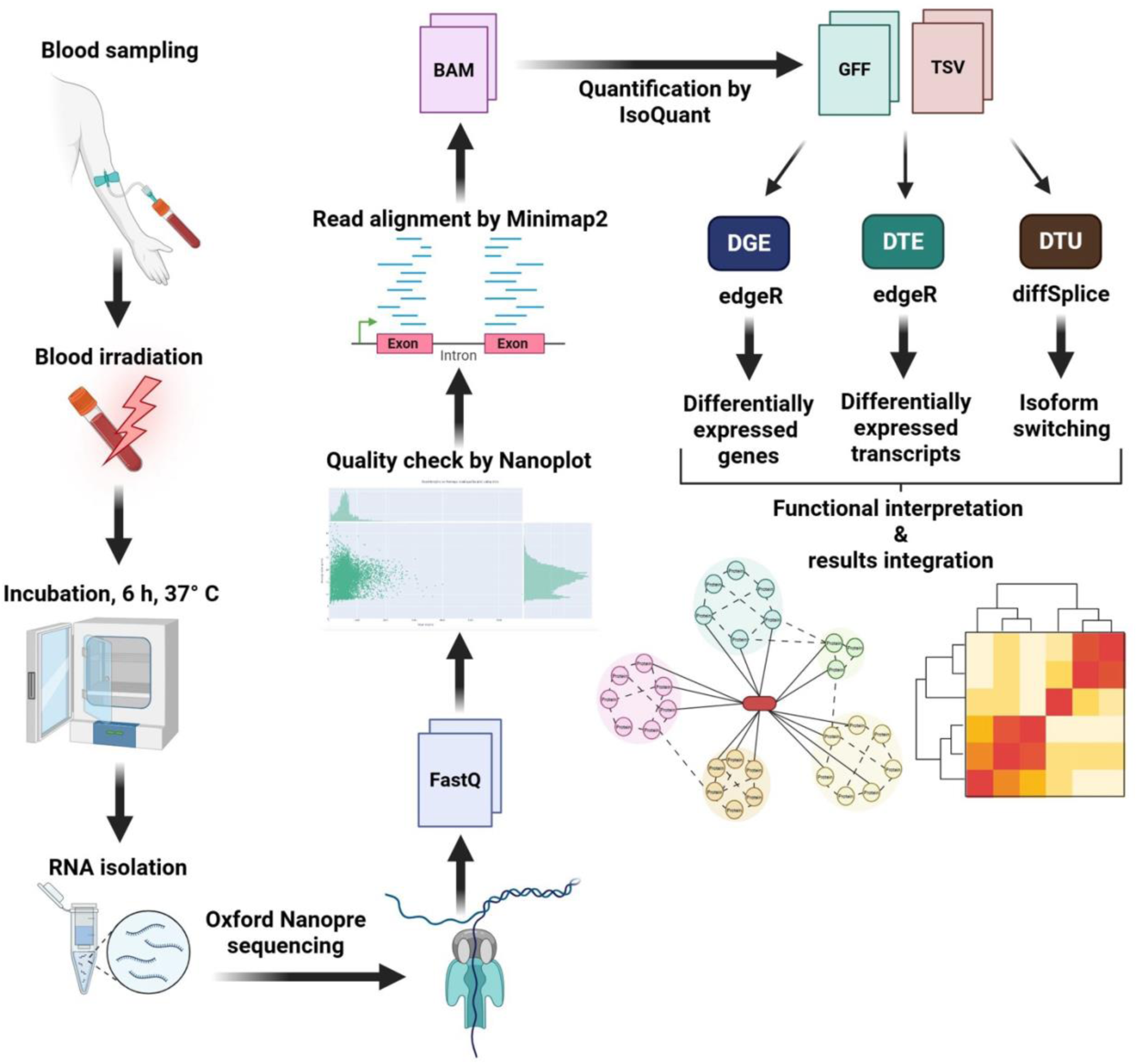
Experimental and analytical workflow of the study. Blood samples were collected from three healthy donors in EDTA tubes, exposed to 4 Gy X-rays, and incubated for 6 h at 37 °C. RNA was isolated from total leukocytes and sequenced using ONT. The quality of the generated FASTQ files was checked using NanoPlot software, and reads were aligned using Minimap2. Generated reads were quantified using the IsoQuant software, and the resulting count matrix was then used for differential gene expression (DGE), differential transcript expression (DTE), and differential transcript usage (DTU) analyses. Created in BioRender. Zahnreich, S. (2026) https://BioRender.com/bf6f6x2.

Exploratory data analysis revealed substantial inter-donor variability in gene expression profiles (Figure 2A), highlighting the importance of accounting for donor-specific effects in the downstream analysis. To select a good-performing analytical framework for modeling our ONT dataset, we compared the performance of two statistical modeling approaches, DESeq2 and edgeR, on identical filtered feature sets at the gene and transcript level. At the gene level (13,020 genes surviving expression filtering), log2 fold-change (LFC) estimates were highly correlated between frameworks (R = 0.99; Figure 2B); edgeR identified 183 significant genes and DESeq2 identified 120 (FDR ≤ 0.05), with 117 genes shared between frameworks, 66 identified only by edgeR, and three identified only by DESeq2 (Figure 2C). At the transcript level (14,146 transcripts), agreement was similarly high (R = 0.99); edgeR identified 119 significant transcripts, and DESeq2 identified 71, with all 71 DESeq2 hits also identified by edgeR (i.e., DESeq2’s significant transcript set was a complete subset of edgeR’s) (Figure 2D). We used edgeR as our primary framework throughout gene-, transcript-, and splicing-level analyses for methodological consistency.

**Figure 2:**
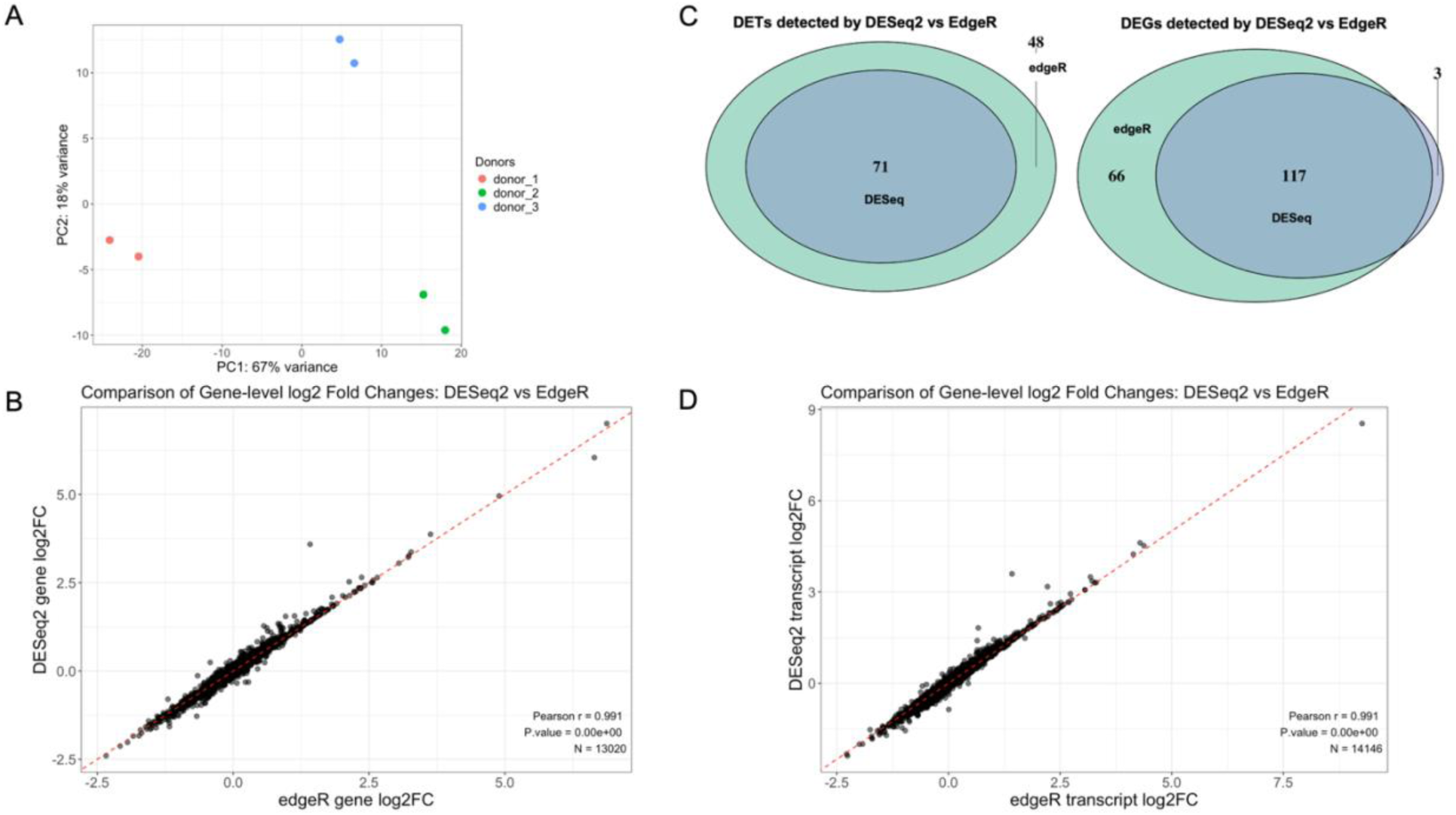
Exploratory ONT long-read data analysis. (A) Principal Component (PC) Analysis plot showing interdonor variation. (B) Dot plot of gene-level log2 fold-changes calculated by both edgeR and DESeq2 statistical frameworks. (C) Venn diagram showing the overlap of differentially expressed transcripts (DETs; left panel) and differentially expressed genes (DEGs, right panel) obtained by edgeR and DESeq2. (D) Dot plot of transcript-level log2 fold-changes calculated by both edgeR and DESeq2 statistical frameworks.

Next, to identify transcriptional changes specifically associated with IR, DEA was performed, using a model that included the donor as a blocking factor to control for individual variability. This analysis identified 66 downregulated and 117 upregulated genes that were significantly differentially expressed upon 4 Gy X-ray exposure (Figure 3A and Additional File 2). Among these, several well-established IR-responsive genes, including *FDXR*, *CDKN1A*, and *CD83*, were prominently upregulated (Figure 3B).

**Figure 3:**
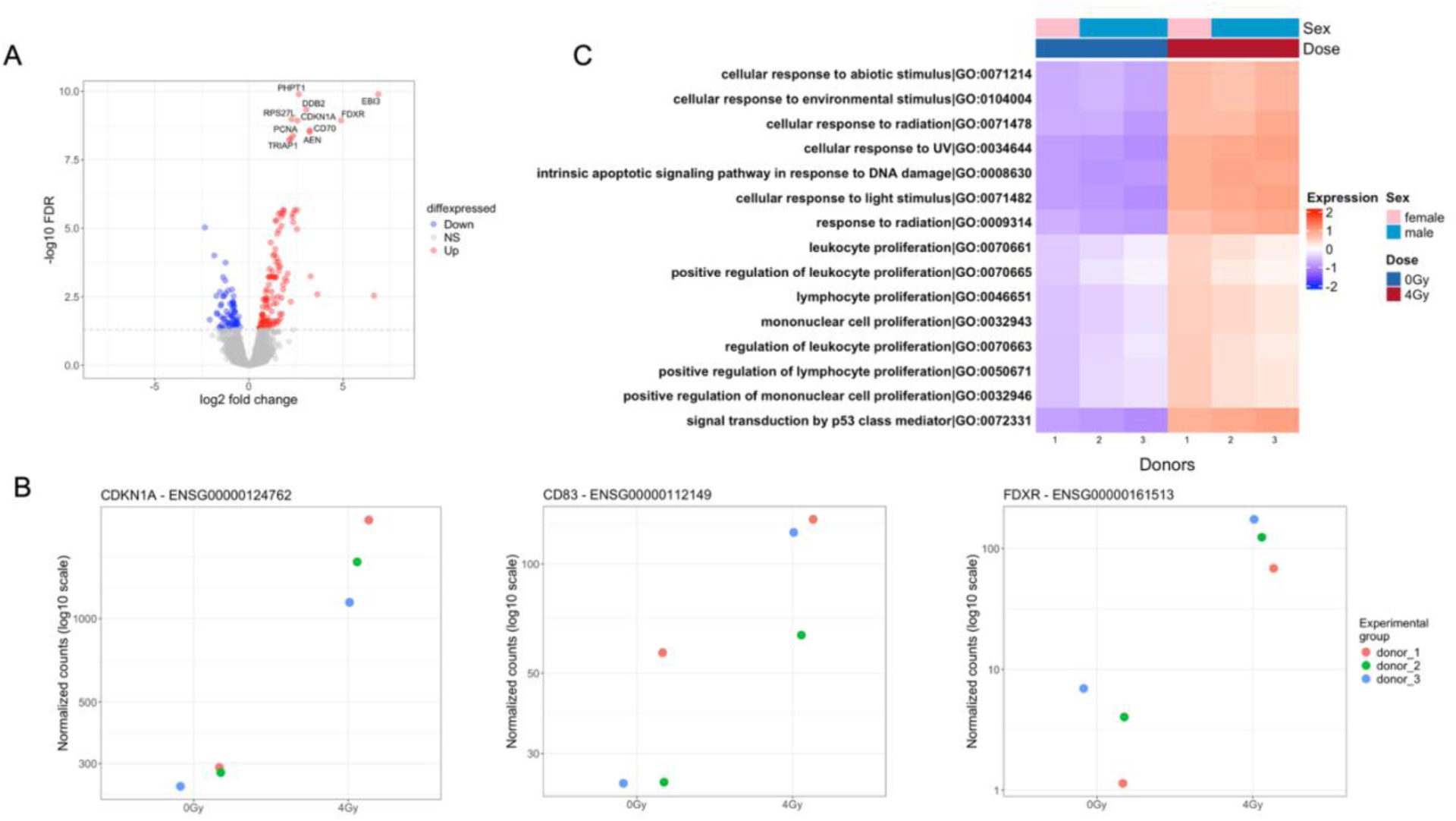
Gene expression and functional enrichment analysis after X-ray exposure. (A) Volcano plot showing the effect of 4 Gy X-rays on gene expression of total leukocytes derived from 3 donors compared to 0 Gy. (B) Gene expression plots of CDKN1A, CD83, and FDXR following X-ray irradiation. (C) Heatmap of the top 15 regulated pathways in response to 4 Gy irradiation compared to 0 Gy.

To place the DEGs in a biological context, we performed functional enrichment analyses using an explicitly defined background universe consisting of the 13,020 genes passing expression filtering. KEGG pathway analysis revealed significant enrichment of the p53 signaling pathway (hsa04115; GeneRatio = 15/89, p_adj_ = 1.8×10^−12^), consistent with activation of DNA damage checkpoints and apoptosis-related mechanisms. Gene Ontology (GO) enrichment analysis of the 183 differentially expressed genes further highlighted B cell proliferation (GO:0042100, p_adj_ = 0.0012) and positive regulation of lymphocyte activation (GO:0051251, p_adj_ = 0.010) among the significantly enriched terms (Figure 3C and Additional File 3).

Additionally, we performed GSEA to gain a more detailed understanding of immune modulation induced by IR. This analysis revealed downregulation of antigen presentation and processing pathways alongside upregulation of inflammatory responses, such as the humoral immune response (Additional Files 4 and 5).

### Transcript-level analysis reveals isoform-specific regulation and novel transcript detection upon irradiation

We further extended our analysis to the transcript level to capture isoform-specific regulation following irradiation. Differential transcript expression (DTE) analysis identified 102 upregulated and 17 downregulated transcripts (Additional File 6). Notably, *IL32* (for which 19 transcript isoforms are annotated in GENCODE v47) was not detected as differentially expressed at the gene level. However, our analyses identified two transcript isoforms that were significantly upregulated: ENST00000440815, with average raw counts of 66 and 140 at 0 and 4 Gy, respectively, and ENST00000325568, with average raw counts of 6 and 24, respectively (LFC 1.19 and 1.99, respectively; Figure 4A). Importantly, leveraging the LR capability of ONT to span full-length transcripts, we detected 26 previously unannotated transcripts out of 22,512 (3,579 passed low-expression filtering (filterByExpr)) constructed by IsoQuant, of which 24 were significantly upregulated, and two were significantly downregulated. These may represent novel isoforms or transcript variants of known genes. As an *a priori* read-support criterion, we required ≥3 assigned reads in at least two of the three donors per condition; all 26 transcripts met this threshold (Additional File 7).

**Figure 4:**
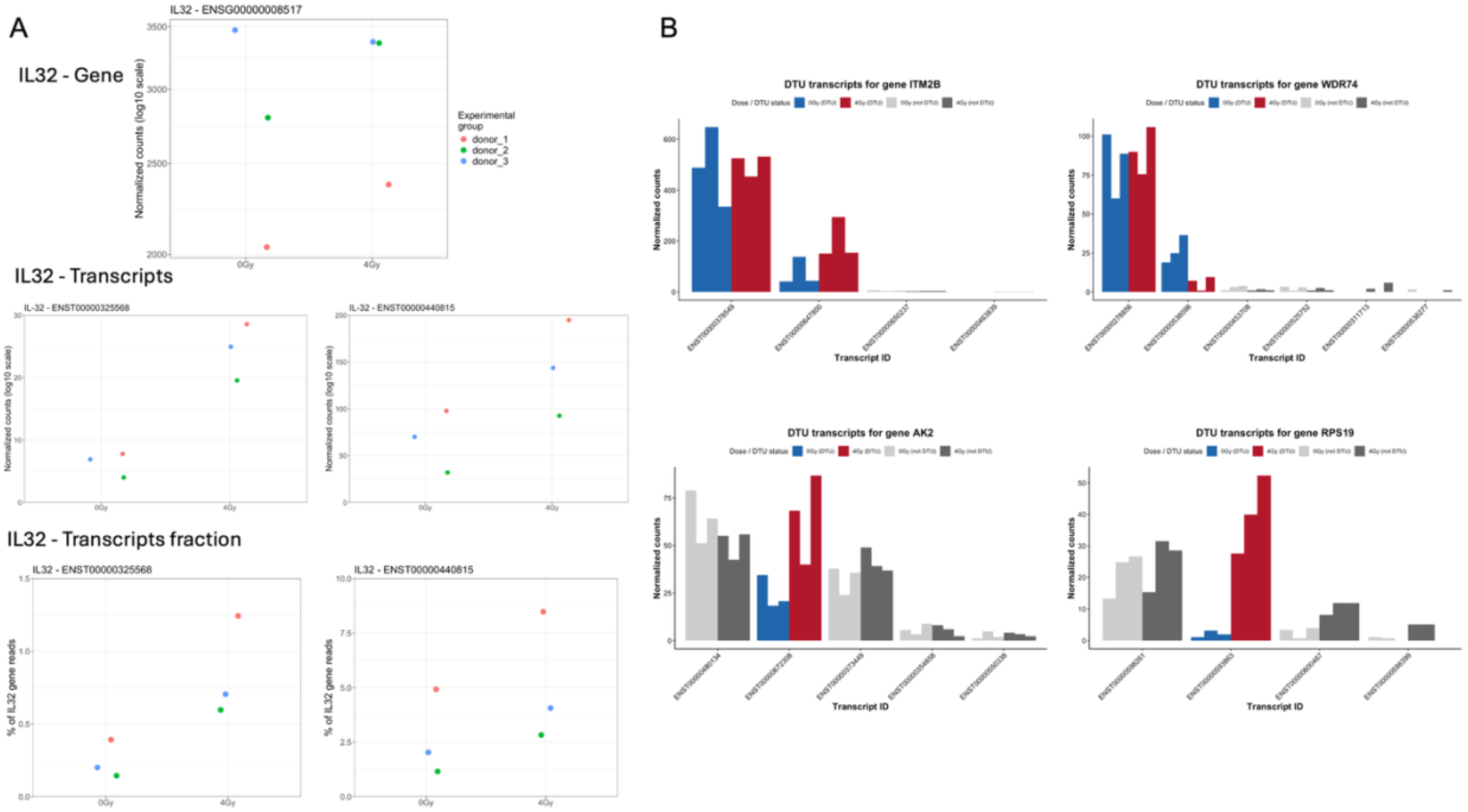
The effect of X-rays on transcript expression and isoform switching. (A) Expression profiles of IL32, showing average gene-level expression across all IL32 isoforms (top panel), normalized expression of the two differentially expressed transcript isoforms (middle panel), and their relative isoform fractions as percentages of total IL32 gene reads (bottom panel). (B) Plot representing transcript isoform distribution of ITM2B, WDR74, AK2, and RPS19 in response to 4 Gy X-rays.

To further explore isoform-level regulation, we performed DTU analysis, which examines changes in the relative abundance of transcript isoforms arising from alternative splicing events. We found that *RPS19*, *WDR74*, *AK2*, and *ITM2B* exhibited IR-associated changes in isoform usage (Figure 4B, Additional File 8). While *ITM2B* was also detected as a DEG, all four genes were also identified in the DTE analysis. Interestingly, although DTE detected only one transcript per gene as significantly regulated, DTU revealed changes in transcript usage for *WDR74* and *ITM2B*, each involving two significant isoforms, highlighting isoform redistribution in response to IR. None of the 26 unannotated transcripts showed significant DTU.

Next, we focused on the 26 previously unannotated transcripts to evaluate protein-coding potential. Using TransDecoder, an ORF was predicted for 11 of the 26 transcripts (14 individual ORFs in total, as three transcripts each had two independently predicted ORFs); the remaining 15 transcripts had no detectable ORF. Of the 11 transcripts with a predicted ORF, eight had at least one complete ORF (containing both start and stop codons), two had only a partial ORF (missing either a start or stop codon), and one had only an internal ORF (lacking both) (Additional File 9). To further assess coding potential, we employed CPC2 and CPAT. Of the 26 transcripts, CPC2 classified 8 as coding (Additional File 10). CPAT could not be evaluated for one transcript (transcript25470.chr1.nic), which contained no ORF detectable by this tool; of the remaining 25 evaluable transcripts, CPAT, using the standard coding-probability threshold of 0.5, classified 11 as coding (Additional File 11). All 8 transcripts classified as coding by CPC2 were also classified as coding by CPAT and had a TransDecoder-predicted ORF of any completeness; we refer to these 8 transcripts as high-confidence coding candidates (Figure 5A, Additional File 12), noting that CPC2’s coding classification is the limiting criterion among the three tools for this set.

**Figure 5:**
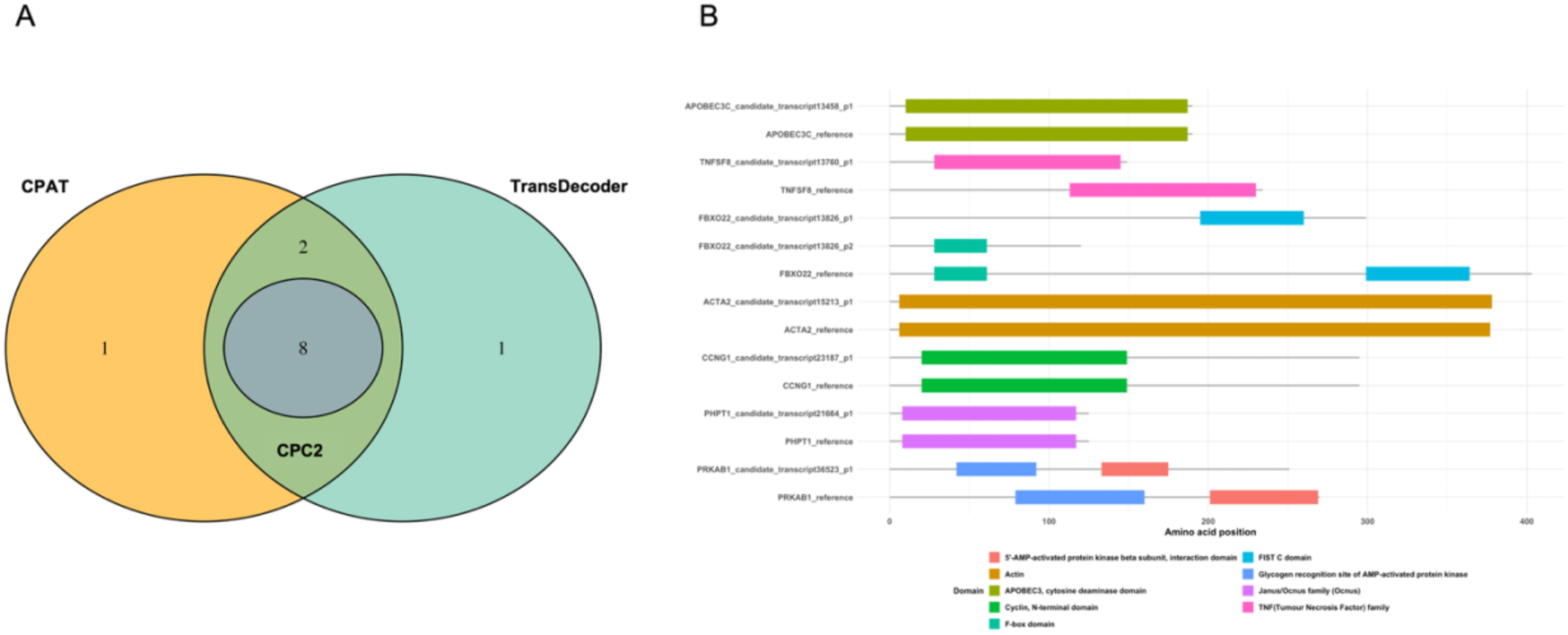
Analysis of unannotated transcripts. (A) Venn diagram of unannotated transcripts with coding potential predicted by CPC2, CPAT, and TransDecoder. (B) Predicted protein domain architecture of the two candidate transcripts (transcript21664, transcript13826) compared to their GENCODE reference proteins.

Annotation analysis using GffCompare showed that these eight transcripts were associated with known gene loci but exhibited structural differences from the corresponding reference annotations. Four transcripts were assigned class code j (transcript13458.chr22.nnic, transcript13760.chr9.nnic, transcript13826.chr15.nnic, and transcript15213.chr10.nnic), representing novel combinations of splice junctions sharing at least one junction with the reference transcript, and were associated with *APOBEC3C*, *TNFSF8*, *FBXO22*, and *ACTA2*. Two transcripts were assigned class code k (transcript21664.chr9.nnic and transcript23187.chr5.nnic), denoting same-strand containment of the reference transcript by the novel model, consistent with a 5′- or 3′-extended isoform, and were associated with *PHPT1* and *CCNG1*. One transcript (transcript36523.chr12.nnic), associated with *PRKAB1*, was assigned class code n, denoting a novel intron chain only partially contained within the reference intron structure. One transcript was assigned class code o (transcript61117.chr1.nnic), denoting same-strand exonic overlap with the reference transcript without a shared splice junction; this transcript has no assigned gene symbol but is annotated to the Ensembl gene ID ENSG00000289589. Extending this classification to all 26 unannotated transcript models identified in this study (Additional File 13), the distribution was: c (query transcript fully contained within a reference transcript) = 3, i (fully contained within a reference intron) = 1, j = 10, k = 4, n = 2, o = 5, and x (exonic overlap with a reference transcript on the opposite strand) = 1.

BLASTp searches of the predicted ORFs against the NCBI nr database confirmed strong homology to the corresponding human reference proteins (*APOBEC3C*, *TNFSF8*, *FBXO22*, *ACTA2*, *PHPT1*, *CCNG1*, and *PRKAB1*) for seven of the eight candidates (≥94% identity, E-values ≤10⁻¹⁷; Additional File 14), supporting that these represent genuine translated products of the assigned genes rather than spurious ORFs. The eighth candidate (transcript61117.chr1.nnic, for which TransDecoder identified only a partial ORF) returned no significant BLASTp hits.

To further evaluate whether the 8 novel potential protein-coding candidates encode altered protein domains, we performed InterProScan analysis on the predicted ORFs by TransDecoder of the 8 candidate genes. Seven of the eight candidate genes had a corresponding GENCODE v47 reference protein available for domain-architecture comparison. Four of eight genes (*ACTA2*, *APOBEC3C*, *CCNG1*, *PHPT1*) showed no domain-level differences between candidate and reference proteins. Two genes (*PRKAB1*, *TNFSF8*) retained their functional domains but with altered position. *FBXO22* showed the clearest evidence of domain-level consequences: two candidate ORFs arising from the same transcript locus each retained only one of the two domains present in the reference protein (F-box domain or FIST C-domain, respectively), suggesting the isoform may disrupt the canonical F-box/FIST domain pairing required for SCF-complex-mediated substrate recruitment (Figure 5B, Additional File 15).

Applying SQANTI3’s standard structural quality filter, 19 of the 26 candidate transcripts passed without an artefact flag. For each of these 19, SR junction support was independently re-derived from STAR SJ.out.tab and LR support was independently confirmed by direct CIGAR inspection of ONT alignments, in both cases at the exact coordinates of each transcript’s defining junction, across all six sequencing libraries (Additional File 16).

Two transcripts, associated with *FBXO22* (transcript13826.chr15.nnic, class code j) and *PHPT1* (transcript21664.chr9.nnic, class k), showed independent coding-potential evidence across four tools (CPC2, CPAT, TransDecoder, SQANTI3/psauron). *PHPT1*’s transcript was flagged by SQANTI3 as spanning an adjacent, GENCODE-annotated overlapping locus (*AJM1*). We examined the expression of *AJM1* in our dataset and found negligible, non-dose-dependent read counts (1–3 reads across all six libraries), inconsistent with *AJM1*-driven transcriptional readthrough and supporting a *PHPT1*-specific novel transcription start site. *PHPT1*’s novel junction was directly confirmed by both SR (36 reads) and LR (present in all six libraries); its six other, already-annotated junctions were also confirmed by both read types. *FBXO22*’s transcript contains two junctions flagged as novel by SQANTI3. One reflects a novel combination of two individually annotated splice sites and was confirmed by both SR (30 reads) and LR (29 reads, present in all six libraries). The other introduces a genuinely novel splice donor site not previously annotated at either individual site; this junction had no SR support and LR support limited to 4 reads across only two of six libraries, both irradiated samples, with none in any control replicate. *FBXO22*’s remaining five already-annotated junctions were likewise extensively confirmed by both read types.

Among the remaining 17 filter-passing transcripts, none showed coding-potential evidence. Two transcripts (transcript30521.chr2.nnic, class o; and transcript26863.chr12.nnic, class k) showed the strongest independent support for their defining novel junction (74 and 70 confirming LRs respectively, both in all six libraries); both loci showed clear SR splicing activity nearby, though not at the exact novel junction itself. A further five transcripts (transcript926.chr3.nnic, class j; transcript31443.chr6.nnic, class o; transcript33002.chr12.nnic, class j; transcript8598.chr15.nnic, class j; transcript13844.chr22.nnic, class o) showed the same qualitative pattern at lower read numbers (4–7 confirming long reads, present in three or four of six libraries). Two transcripts (transcript25470.chr1.nic, class j, and transcript34121.chr1.nic, class c) showed SR and strong LR confirmation of their full junction structure. One transcript (transcript26870.chr12.nnic, class k) was likely a minor (∼40 bp) shift from an already-supported SR junction. Two transcripts (transcript61117.chr1.nnic and transcript61119.chr1.nnic, both class o) showed no meaningful SR activity anywhere in the surrounding locus; LR support for these two was itself inconsistent (one read in a single library for one transcript; 17 reads in five of six libraries for the other). One further transcript (transcript39964.chr19.nnic, class x) showed SR splicing activity nearby, but only one LR supporting the novel junction, in one library. Four transcripts (transcript20940.chr17.nnic, class c; transcript26885.chr12.nnic, class i; transcript29638.chr6.nnic, class n; transcript32230.chr3.nnic, class c) were mono-exon and could not be evaluated by this junction-based approach.

Taken together, this analysis identified candidate novel isoforms encompassing both protein-coding and non-coding candidates. Among the strongest candidates, some showed cross-platform support for their novel junctions (transcript25470.chr1.nic, transcript21664.chr9.nnic, and transcript13826.chr15.nnic). Others were confirmed by exact CIGAR-matched LR support but were not detected in the SR data (transcript30521.chr2.nnic and transcript26863.chr12.nnic), potentially reflecting the greater sensitivity of LR sequencing for isoform-specific regulation that may fall below the detection threshold of standard SR approaches.

## Discussion

The present study represents the first systematic DTU application of ONT LR RNA-seq for transcriptomic profiling of leukocytes from irradiated human whole blood, enabling the analysis of IR-induced expression changes at both the gene and transcript levels. We identified 183 genes and 119 transcripts that were significantly regulated in leukocytes from whole blood exposed to 4 Gy X-rays, including candidate unannotated transcript models, some of which come from known radiosensitive genes. These findings provide evidence of IR-induced isoform switching, which may lead to alterations in protein structure and function. We used blood as a model to investigate radiation effects because blood is a readily accessible and representative system for studying biological responses to *ex vivo* or *in vivo* IR exposure.

To assess whether the reported gene-level findings were driven by a single donor, we performed a leave-one-donor-out sensitivity analysis using the same edgeR framework. 76 to 124 of the 183 original DEGs (42–68%) remained significant, reflecting the expected loss of power at n = 2. However, LFC estimates remained highly correlated with the full three-donor model after removing any single donor (Spearman r = 0.80–0.86), which is a more direct indicator of donor-independent consistency than the raw DEG retention count alone.

At the transcriptional level, many studies have examined IR-related gene expression in human whole blood or isolated peripheral blood mononuclear cells (PBMCs) using microarrays or targeted qPCR [26, 69–71]. For example, the meta-analysis by Biolatti et al. on five microarray datasets identified 21 genes that were differentially expressed in response to 0.5, 1, 2, 3, 4, 5, and 6 Gy of low LET IR at 24 h post-irradiation, including *BAX*, *FDXR*, *DDB2*, *PCNA*, and *GADD45A* [72]. But so far, only a few studies, including ours, have conducted genome-wide transcriptome analyses with RNA-seq [17, 40, 73, 74]. Among these, only a study by Cruz-Garcia et al. with isolated PBMCs used ONT LR RNA-seq [40]; all others relied on SR data. However, they primarily focused on validating established radiation-induced gene signatures using the ONT LR technique for on-site biodosimetry. But LR offers strong potential for biodosimetric and analytical applications by providing isoform-level resolution, reducing ambiguity in isoform reconstruction, and improving detection of splice variants alongside gene expression, enabling deeper insight into cellular IR responses. IR has already been found to modulate alternative transcription and splicing of specific genes [75, 76]. Developing a high-resolution, highly specific IR signature could yield valuable insights into exposure scenarios, including radiation type or mixed fields, homogeneity (partial-versus total-body), and dose rate.

We observed upregulation of the DDR and inflammatory genes, including *EBI3*, *FDXR*, *DDB2*, *BAX*, *CCR7*, *TNFAIP6*, and *CD83*. *LNCTAM34A*, a long non-coding RNA (lncRNA), was also found to be upregulated. It is known to be induced by *TP53* and positively regulates miR34a expression, causing G1 arrest [77, 78]. Collectively, these results indicate that X-ray irradiation triggers a coordinated transcriptional program encompassing DDR and immune modulation in human leukocytes. Importantly, our ONT dataset successfully recapitulated several well-established IR-responsive genes, which are known markers of DNA damage and p53 pathway activation [26, 79, 80]. This concordance provides strong validation for the accuracy and biological relevance of ONT-based transcriptomic profiling in the context of biodosimetry.

Interestingly, two transcript isoforms of *IL32* were identified as differentially expressed transcripts (DETs) despite the absence of a corresponding significant gene-level change (DTE without DGE), whereas two isoforms of *WDR74*, one of which was identified as a DET, exhibited significant DTU without a corresponding gene-level change. In contrast, *ITM2B* showed significant changes at both the DTU and gene-levels, with one transcript isoform was detected as DET.

To evaluate whether the transcript-level findings identified here are uniquely enabled by LR sequencing, we compared the LR dataset with our previously generated SR dataset obtained from the same donors across a broader range of doses (0.5-4 Gy) and time points (2 and 6 h post-exposure) [17]. However, for the comparison, we restricted the analysis to the same two dose groups (0 and 4 Gy) and time point (6 h) used in the LR study. The SR dataset served as a reference dataset owing to its higher per-base sequencing accuracy and matched irradiation conditions, time point, and donors relative to the LR dataset. We performed matched DTE and DTU analyses on the SR dataset, applying identical significance procedures (edgeR for DTE; the diffSplice/stageR two-stage procedure for DTU, both after applying filterByExpr) to both datasets. Of the two differentially expressed *IL32* transcripts identified in the LR data, one (ENST00000440815) did not pass expression filtering in the SR dataset and could therefore not be tested, while the other (ENST00000325568) was tested but did not reach significance (FDR = 0.90), together suggesting that this finding is not readily accessible to SR sequencing at comparable sample size. For the DTU analysis, no gene reached significance in the SR DTU analysis under the matched stageR procedure at a 5% overall FDR. This highlights the enhanced capability of LR sequencing to capture isoform-specific regulatory events that are often masked in conventional gene-level analyses by SR RNA-seq.

Comparison of gene detection between our LR (IsoQuant, GENCODE v47) and SR (Salmon/tximeta, GENCODE v46) datasets requires a common gene universe and an identical detection threshold, since the two pipelines differ in quantification method and annotation version, in addition to sequencing platform. Restricting both datasets to the shared set of annotated genes and applying the same low-expression filter (filterByExpr) to each, the SR dataset detected more genes overall (12,329 vs. 10,650). However, 1,470 genes detected in the LR dataset were not detected in the SR dataset; these were disproportionately non-coding, comprising lncRNAs (504/1,470) and processed pseudogenes (574/1,470), with protein-coding genes representing only 19% (273/1,470) of this set. Conversely, 3,149 genes were detected only in the SR dataset.

We additionally compared DEGs identified from our SR and LR datasets, restricting the SR data to the same two dose groups and time point (0 Gy and 4 Gy, 6 h, n = 3 donors each) used in the LR study, rather than the full dose series used in our original published analysis of this dataset; DEG counts reported here for the SR dataset therefore reflect this restricted two-group comparison and may differ from those reported in the original publication. DESeq2 was used for the SR analysis with independent filtering, consistent with our previously published methodology for this dataset; edgeR with filterByExpr-based filtering was used for the LR analysis, consistent with the primary DEA presented above. Of 116 significant genes (FDR ≤ 0.05) identified in the SR dataset and 183 identified in the LR dataset, 58 were shared between platforms, 58 were significant only in the SR dataset, and 125 were significant only in the LR dataset (Additional File 18).

Additionally, we extended our analysis to compare a set of 34 genes that we previously reported as new radiosensitive genes in our SR study. Of these 34 genes, 29 were present in the LR gene-level results after filtering, while all 34 were present in the full SR differential expression results (also restricted to the same two dose groups (0 and 4 Gy) and time point (6 h) used in the LR study, with DESeq2 independent filtering applied as in the original publication; Additional File 19). Only one gene (*G0S2*) reached significance (FDR ≤ 0.05) in the LR dataset. Considering direction of effect rather than significance alone, 18 of the 29 genes with values available in both datasets showed concordant directions of change between platforms (62.1%); this was not significantly different from the 50% expected by chance (binomial test, P = 0.265). However, comparison of the continuous effect-size estimates showed a significant positive association between LR and SR LFC (Spearman’s ρ = 0.41, P = 0.027), indicating moderate agreement in the relative magnitude and direction of gene-level responses between platforms. A gene-set-level test (mroast, limma) treating the 34-gene signature as a single set within the LR differential expression model showed a nominal trend toward differential expression in the mixed-direction test (P = 0.081). Together, these findings provide some evidence of consistency with the previously identified radioresponsive signature, but do not provide sufficient statistical support for full replication in the LR dataset. However, as a broader assessment of cross-platform concordance, we also tested the complete set of 116 SR DEGs as a predefined gene set in the LR dataset. This showed strong evidence of a coordinated response (mixed-direction test, *P* < 0.0005), driven predominantly by the SR-upregulated genes. These discrepancy between the SR and LR datasets may reflect limited statistical power given the small sample size (n = 3) and the single dose and time point examined, as well as genuine differences between both dataset, including differences in blood processing and RNA extraction methods, platform-specific biases, read-to-transcript assignment ambiguity, and quantification sensitivity across different pipelines, precluding a fully controlled comparison of platform performance alone, which were also reported in other studies [81–83]. Despite these differences in the specific DEG sets identified, both datasets recovered a broadly consistent overall biological response, including activation of DNA damage and immune-related pathways, supporting the general validity of the LR transcriptomic signal while underscoring that platform-, pipeline-, and framework-level differences meaningfully affect the specific gene sets recovered.

Integrating structural quality control with our coding-potential analyses substantially refines which of the 26 candidate transcripts warrant higher confidence. *FBXO22* and *PHPT1* emerge as our strongest candidates of the potential coding unannotated transcript models, showing convergent evidence across four independent coding-potential tools, a genuine novel splice junction, and no unresolved artefact flags. Among the remaining transcripts, a subset showing no coding potential but a genuinely novel, multi-exon splice structure represents plausible candidate lncRNAs. Several candidate non-coding transcripts identified here map to genes with established relevance to radiation and hematopoietic biology, including *MDM2* (transcript26863.chr12.nnic), a central regulator of the p53 DNA-damage response, *TAL1* (transcript25470.chr1.nic), the master transcription factor of hematopoiesis, and *MZT2B* (transcript30521.chr2.nnic). Two other transcripts, *APOBEC3C* (transcript13844.chr22.nnic) and *IER5* (transcript61119.chr1.nnic), showed comparatively lesser independent read-level support and should be interpreted with additional caution relative to the candidates above. While coding-potential prediction did not support these as protein-coding isoforms, their genomic location overlaps these genes’ loci and might have regulatory functions over those genes. Other non-coding candidates were associated with already known lncRNAs, including *THUMPD3-AS*1 (transcript926.chr3.nnic) and *VAV3-AS1* (transcript34121.chr1.nic). Together, this analysis identifies a set of candidate isoform models varying in confidence, with the strongest convergent evidence for genuine novelty found in transcript13826.chr15.nnic (*FBXO22*) and transcript21664.chr9.nnic (*PHPT1*) among the coding candidates, and transcript30521.chr2.nnic, transcript25470.chr1.nic, and transcript26863.chr12.nnic among the non-coding candidates. Overall, while our analyses provide evidence of novel isoform discovery and open avenues for better understanding radiation-induced transcriptomic events, orthogonal experimental validation such as targeted reverse qPCR or Sanger sequencing across the novel junctions would be required to confirm the transcript structures and biological functions proposed here, particularly for the lower-confidence candidates.

To assess the effect of sequencing depth on differential expression detection, we performed a read-depth saturation analysis at the gene-level. The filtered gene-level count matrix was subsampled to 5-100% of the full sequencing depth, with 20 independent replicates at each depth, and the same edgeR quasi-likelihood model, including donor as a blocking factor, was refitted for each subsample. Among the 183 genes identified as significant at full depth, the mean recovery increased from 7.8% at 5% depth to 29.2% at 10%, 54.2% at 30%, 70.1% at 50%, 86.7% at 80%, and 93.3% at 90% depth. Genes with larger absolute effect sizes were generally detected more reliably at lower sequencing depths. Overall, 28 of 183 genes (15.3%) showed robust detection across subsampled depths, including known radiosensitive genes (*CDKN1A*, *FDXR*, *AEN*, *MDM2*), whereas 67 (36.6%) were classified as depth-sensitive. In parallel, agreement with the full-depth effect estimates increased with sequencing depth, with mean Spearman correlation rising from 0.364 at 5% depth to 0.838 at 50%, 0.949 at 80%, and 0.976 at 90%; mean concordance increased from 0.236 to 0.872, 0.965, and 0.984, respectively. The total number of significant genes increased from a mean of 14 at 5% depth to 137 at 50% and 182 at 90%, compared with 183 at full depth. Together, these results indicate that differential expression detection remains partly limited by sequencing depth in this LR dataset, with approximately 90% of the current sequencing depth required to recover >90% of the full-depth DEG set, while effect-size estimates were already highly concordant with the full-depth analysis at this depth (Additional File 20). Collectively, these results suggest that detection sensitivity is approaching saturation at current sequencing depth, with further gains expected to be modest and concentrated among lower-effect-size transcripts.

A limitation of this study is the small number of biological replicates (n = 3 per group) with a single dose and time point, which might constrain the statistical power available for both differential expression and differential transcript usage analyses. Irradiation was performed *ex vivo* on whole blood, which does not fully recapitulate the physiological conditions of in vivo exposure. The findings provided in this study would benefit from orthogonal experimental confirmation and validation in a larger, sex- and age-balanced cohort. Additionally, as no differential blood count or computational cell-type deconvolution was performed, we cannot distinguish compositional shifts in cell-type proportions from genuine per-cell transcriptional changes in the observed differential expression signature.

Although these approaches are mostly focused on hypothesis generation for the role of these identified transcripts, they can be a valuable toolkit to elucidate unknown and novel roles for transcriptomic features. We believe that the workflow we propose in this study can be adapted to existing analysis pipelines for other experimental settings. Overall, we highlight the potential of LR RNA-seq for biodosimetry and radiation biology applications, and provide a practical pipeline to encourage broader adoption of this technology in the field, which, in combination with classical SR transcriptomics, could deliver a more complete characterization of radiation-induced transcriptional responses.

## Conclusion

This study demonstrates that radiation exposure is associated with transcriptomic changes in human leukocytes at the isoform level, including differential transcript expression and usage in genes such as *WDR74*, *AK2*, *ITM2B*, and *RPS19*, and candidate novel protein-coding transcripts, including *FBXO22* and *PHPT1*, and other long non-coding candidate isoforms. Notably, *FBXO22* showed evidence of altered protein domain architecture relative to the reference isoform. Using LR RNA-seq, we confirmed known radiation response signatures at the gene level and identified transcript-level events that were not detected, or were not testable, in a matched SR dataset from the same donors, suggesting that LR sequencing can capture aspects of the radiation transcriptomic response that are difficult to access with standard SR approaches. Despite the limited sample size, these results illustrate the potential of LR sequencing to contribute additional, isoform-resolved information to studies of the cellular radiation response. Building on these findings, future studies should leverage LR RNA-seq with larger cohorts and additional dose and time points to develop robust, isoform-level biomarkers for biodosimetry and personalized radiation risk assessment.

## Supporting information

Additional File 1: Globin and ribosomal RNA fractions

Additional File 2: Differentially expressed genes

Additional File 3: Enrichment analysis

Additional File 4: Gene set enrichment analysis

Additional File 6: Differentially expressed transcripts

Additional File 7: Unannotated differentially expressed transcript counts

Additional File 8: Differential transcript usage results

Additional File 9: TransDecoder results

Additional File 10: Coding Potential Calculator 2 results

Additional File 11: Coding-Potential Assessment Tool results

Additional File 12: Coding potential summary

Additional File 13: GffCompare

Additional File 14: Blastp

Additional File 15: InterProScan

Additional File 16: SQANTI3

Additional File 17: Leave one donor out differentially expressed genes

Additional File 18: Long and short read differentially expressed genes

Additional File 19: Short vs. long read: 34 differentially expressed genes

Additional File 20: Subsampling analysis

Additional File 5: Gene set enrichment analysis plot

## List of abbreviations

IR: Ionizing radiation
DSBs: Double-strand breaks
LET: Linear energy transfer
RBE: Relative biological effectiveness
PBLs: Peripheral blood lymphocytes
PBMCs: Peripheral blood mononuclear cells
RT: Radiotherapy
DDR: DNA damage response
qPCR: Quantitative PCR
SR: Short-read
LR: Long-read
ONT: Oxford Nanopore Technologies
rRNA: ribosomal RNA
DEGs: Differentially expressed genes
GO: Gene Ontology
GSEA: Gene set enrichment analysis
FDR: False discovery rate
DEA: Differential expression analysis
DTE: Differential transcript expression
DETs: Differentially Expressed Transcripts
DTU: Differential transcript usage
LFC: log2 fold-change
lncRNA: long non-coding
RNA RNA-seq: RNA sequencing
KEGG: Kyoto Encyclopedia of Genes and Genomes
ORF: Open Reading Frame
DGE: Differential Gene Expression
RQN: RNA Quality Number
EDTA: Ethylenediaminetetraacetic acid
CPC2: Coding Potential Calculator 2
CPAT: Coding-Potential Assessment Tool
SQANTI3: Structural and Quality Annotation of Novel Transcript Isoforms
CIGAR: Compact Idiosyncratic Gapped Alignment Report

## Declarations

### Ethics approval and consent to participate

Written informed consent was obtained from all participants before enrolment. The study was conducted in accordance with the Declaration of Helsinki and approved by the Ethics Committee of the Rhineland-Palatinate Chamber of Physicians [No. 2023-17191].

### Consent for publication

Not applicable. The manuscript does not contain data from any identifiable individual.

### Availability of data and materials

Scripts for the analyses presented in the manuscript are available a GitHub - AhmedSAHassan/Phybion_LongReads: ONT pilot for X-ray irradiated samples · GitHub. This repository includes a Conda environment definition (RNAseq_analysis.yaml) specifying software versions, shell scripts for alignment and quantification, and one R Markdown file (PhyBioN_LR_Analysis.Rmd, run in R v4.5.1) containing the complete downstream statistical analysis workflow. The processed datasets, at the transcript- and gene-level, generated and analysed during the current study are available in the GitHub repository https://github.com/AhmedSAHassan/Phybion_LongReads as well as in the Zenodo repository (https://doi.org/10.5281/zenodo.22013472). The individual-level RNA-seq data underlying this article cannot be deposited in a public repository because they contain sensitive human information that is subject to data-protection regulations. In order to safeguard the participants’ privacy and to keep the data pseudonymized, the raw datasets are available only upon reasonable request to the corresponding author.

## Authors’ contributions

Conceptualization: SZ, FM. Data curation: AS. Formal analysis: AS, FM. Funding acquisition: SZ, FM, HS. Investigation: AS. Methodology: FM, AS, SZ. Project administration: SZ. Resources: SZ, FM, HS. Supervision: FM, SZ. Validation: AS, FM, SZ. Visualization: AS, FM, SZ. Writing – original draft: AS, SZ, FM. Writing – review & editing: AS, FM, SZ, HS.

## Acknowledgements

We would like to thank the team (Dr. Christiane Kraemer and Dr. Lukas Hellmann) of the Nucleic Acid Core Facility (NACF) of the Faculty of Biology at the Johannes Gutenberg University of Mainz for their excellent support in sequencing the samples included in this work.

## Additional files

Additional File 1: Globin and ribosomal RNA fractions

Additional File 2: Differentially expressed genes

Additional File 3: Enrichment analysis

Additional File 4: Gene set enrichment analysis

Additional File 5: Gene set enrichment analysis plot

Additional File 6: Differentially expressed transcripts

Additional File 7: Unannotated differentially expressed transcript counts

Additional File 8: Differential transcript usage results

Additional File 9: TransDecoder results

Additional File 10: Coding Potential Calculator 2 results

Additional File 11: Coding-Potential Assessment Tool results

Additional File 12: Coding potential summary

Additional File 13: GffCompare

Additional File 14: Blastp

Additional File 15: InterProScan

Additional File 16: SQANTI3

Additional File 17: Leave one donor out differentially expressed genes

Additional File 18: Long and short read differentially expressed genes

Additional File 19: Short vs. long read: 34 differentially expressed genes

Additional File 20: Subsampling analysis

