## Supplementary figures and images for "Decoding Radiation-Induced Transcriptomic Signatures of Human Leukocytes using Long-Read RNA-Seq: Clinical and Biodosimetric Implications"

### Additional File 5: Gene set enrichment analysis plot

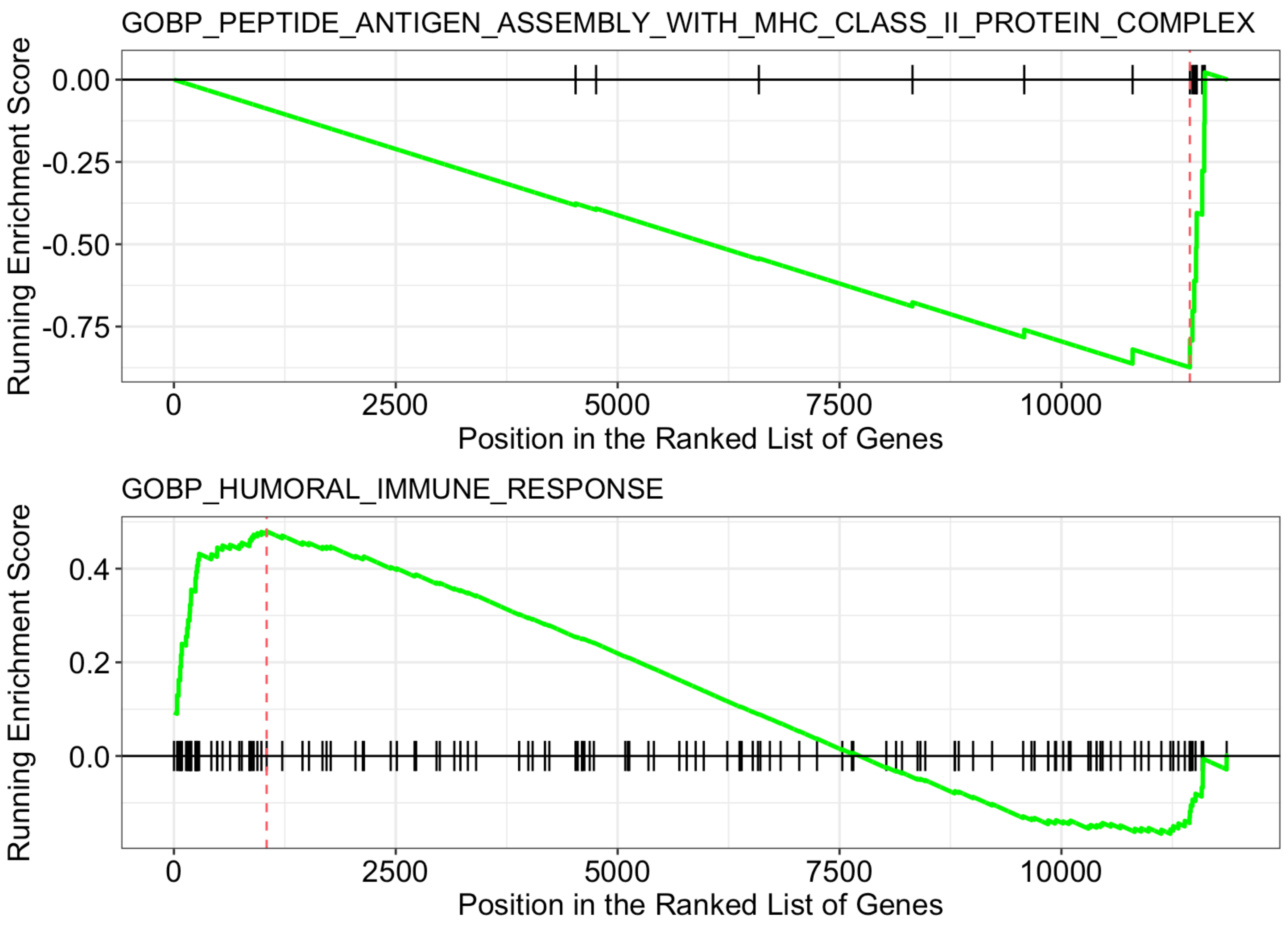
